# Carabid beetle communities along a gradient of agroecological systems: disentangling the roles of pesticides and soil tillage

**DOI:** 10.64898/2026.10.04.756545

**Authors:** Boeraeve Fanny, Noël Grégoire, Bortels Jeannine, Francis Frédéric, Desgain Marie, Sirami Clélia, Vialatte Aude, Caro Gaël, Dufrêne Marc

**Affiliations:** Plant Sciences Axis, Terra Research and Teaching Center, Gembloux Agro-Bio Tech, University of Liege, Belgium; Department of Wildlife Ecology and Conservation, University of Florida, Gainesville, Florida, USA; Entomology, Plant Pathology and Innovative Production Axis, Terra Research and Teaching Center, Gembloux Agro-Bio Tech, University of Liege, Belgium; Biodiversity, Ecosystems and Landscapes Axis, Terra Research and Teaching Center, Gembloux Agro-Bio Tech, University of Liege, Belgium; DYNAFOR Research Unit, UMR 1201, INRAE, Toulouse, France; Laboratory of Agronomy and Environment, UMR1121 – Team of Sustainable Agriculture, Université de Lorraine, Nancy, France

**Keywords:** Agroecology, Carabidae, Farming practices, Organic agriculture, Conservation Agriculture, Community ecology

## Abstract

The potential of alternative agricultural systems, such as organic or conservation agriculture, for biodiversity restoration has been widely investigated. However, variability of practices within each of these two categories of agricultural system has rarely been accounted for. Furthermore, these systems have most often been studied separately, isolating either organic or conservation agriculture (itself often reduced to a soil tillage reduction), and rarely analyzing these systems side by side. This study attempts to disentangle these multiple interrelationships by sampling carabid beetles in a network of 48 commercial Belgian farms including organic agriculture (OA), conservation agriculture (CA), farms combining the two approaches (OCA) and farms applying conventional practices (CVA). In each farm, we sampled carabids using pitfall and emergence traps in one winter cereal field and collected a detailed description of crop management practices. We found that community composition was primarily structured by seasonality early in the season (March–April) followed by convergence (May-June) driven by the aggregation of the most dominant species. Based on agroecosystem classification, organic certification (i.e., the absence of synthetic pesticide use) exerted a twofold stronger effect than application of conservation agriculture principles (i.e., reduced or no soil tillage) on carabid beetle community composition. Notably, OCA supported carabid assemblages that were structurally distinct from those of CA and CVA. Furthermore, carabid communities associated with OA differed significantly in composition from those of CVA. Observed and estimated species richness were also significantly higher in OCA than in CA and CVA, while OA differed from CVA only for observed richness. Beyond differences among agricultural systems, community responses were associated with specific management practices, which explained nearly a quarter of the compositional variation with significant influence of the duration since organic conversion and the frequency of synthetic pesticide application. Carabid abundances were negatively affected by synthetic pesticides and moldboard ploughing, whereas richness was more strongly related to the time since conversion to OA, cover crops, and organic amendments. Overall, these results position synthetic inputs and time since conversion as the main filters over carabid communities. Our study highlights the importance of jointly considering system level analyses and detailed management practices to better understand the complex relationships between agricultural practices and carabid communities.

## Introduction

Intensive agriculture, as practiced in most parts of western countries, has been putting biodiversity under pressure for the last decades. Many studies have been carried out in the last decades on the links between alternative agricultural systems and their impacts on biodiversity^1–3^. Such research ought to guide decision making towards practices less harmful to biodiversity and the environment.

Much of this research concentrates on specific taxonomic groups as ‘bioindicators’ to report on the status of the overall biodiversity. Carabid beetles have long been used, since they are sensitive to disturbances in the environment and are thus indicators of habitat alteration levels^4^, and one of the most worthwhile model groups for biological studies^5^. What’s more, they play a crucial role in agroecosystems by regulating many pests such as aphids, slugs and weed seeds^6,7^. Hence, understanding how agricultural practices impact their community structure is crucial to design crop systems supporting overall biodiversity but also functional biodiversity playing a role in pest control to allow getting rid of synthetic pesticides.

Most scientific work investigating links between agricultural practices and carabids has focused on organic agriculture (OA). OA is probably the most institutionalized alternative agricultural system thanks to its certification scheme recognized worldwide^8^, which forbids all mineral fertilizers and synthetic pesticides. As pesticides represent a direct and indirect mortality cause for a large panel of living organisms in agroecosystems, they act as strong ecological filters influencing the structure of carabid communities. Hence, OA, which gets rid of this filter (as well as other synthetic inputs), usually leads to restored carabid abundance and diversity^1,9,10^. However, because weeds are managed without chemical herbicides, tillage appears to be a very frequent practice in organic agriculture.

Another increasingly popular alternative agricultural system focuses on soil health, as done by the movement of ‘conservation agriculture’ (CA; sometimes also referred to as ‘regenerative agriculture’). CA is based on three principles: reduced tillage without soil inversion, use of cover crops between cash crops, and crop diversification through rotation or intercropping^11^. In practice, however, reduced tillage frequently emerges as the main (or sole) objective, and in any case, the use of synthetic or mineral inputs is not prohibited. Noticeably less studied than OA, probably because less clearly defined and not institutionalized through a legal certification scheme, CA still benefits from a substantial body of literature analyzing impacts of reduced tillage on carabids in comparison with conventional agriculture (CVA). As most carabids species realize their reproduction and juvenile development stages within soils, CA, which gets rid of the ecological filter due to intensive tillage, usually has positive effects on carabid abundance and diversity^12–15^.

Hence, OA and CA both remove two distinct ecological filters of CVA (Figure 1). OA and CA also represent two distinct communities of farmers, each convinced that their approach is the most beneficial to meet current agricultural challenges of producing food while supporting biodiversity. This has created debates on which filter, between synthetic inputs use (in particular, synthetic pesticide use), and intensive soil tillage, impacts biodiversity the most. More recently, some farmers have started to suggest that eliminating both filters would be the panacea with regards to biodiversity^16,17^. However, combining OA and CA (OCA systems) requires a high level of functional biodiversity to ensure natural regulation of pests in the absence of synthetic input use and intensive tillage. While OCA is gaining momentum in the agricultural^18,19^ and scientific arenas^20^, few studies have addressed its potential to restore crucial taxonomic groups such as carabids. In a previous study, Boeraeve *et al.* (2022)^21^ compared OCA and CVA in terms of their carabid and spider communities and showed that OCA restored alpha and beta species diversity in both groups. However, this research was restricted to farming system level comparisons (OCA vs CVA) and lacked a deeper understanding of the gradient of farming systems between OCA and CVA, as well as on the role of detailed practices (timing and intensity of each intervention). Indeed, each system type encompasses a wide range of practices, so that farming systems are better represented as a continuum of practice intensity rather than sharply contrasted categories^9,22^.

**Figure 1:**
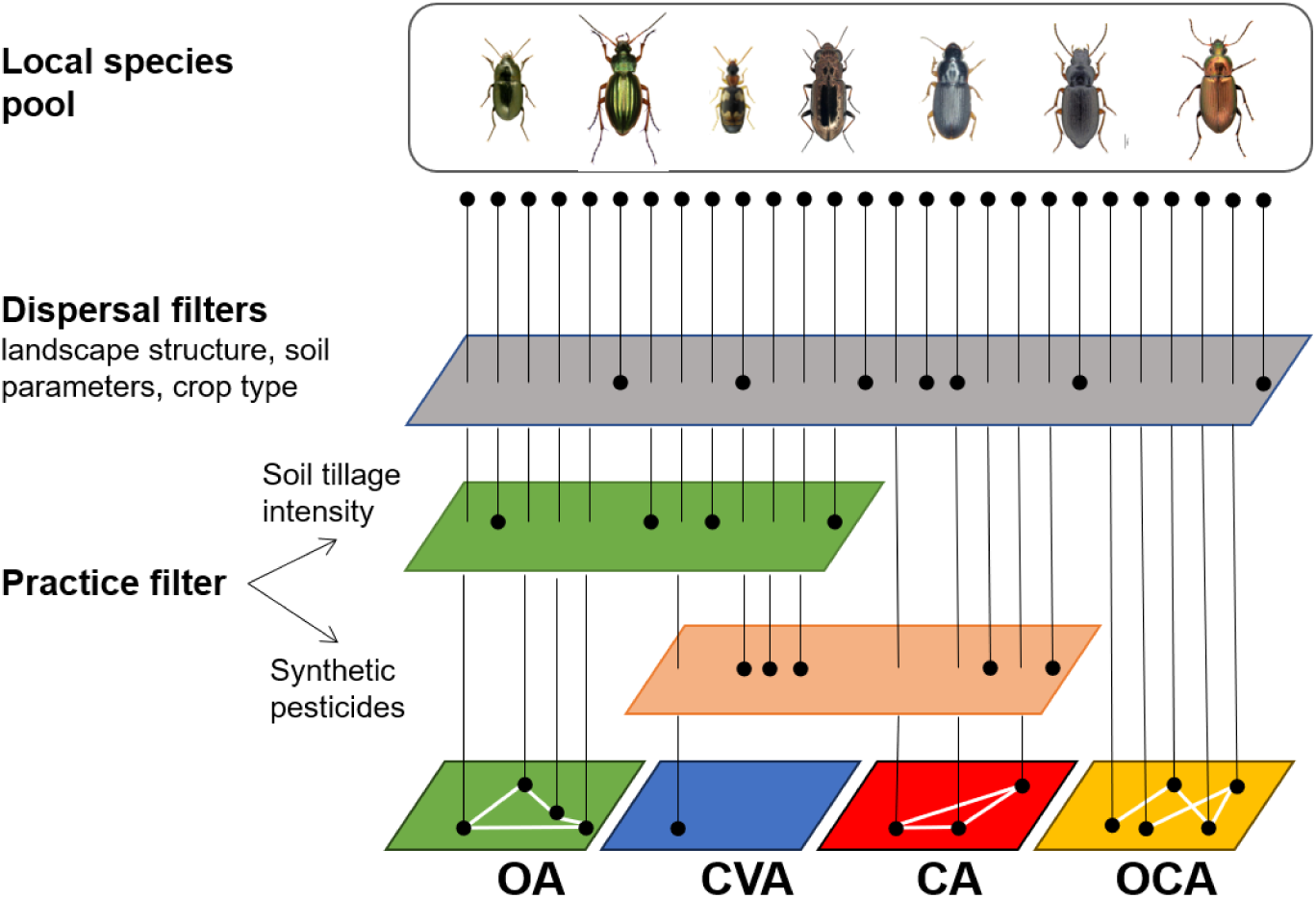
Hypothetical outcomes of organic (OA), conventional (CVA), conservation (CA), and organic-conservation (OCA) agriculture on a local species pool of carabids based on the filter theory (inspired from Lake et al. 2007). The practice filter of synthetic input use (including pesticide) is expected to represent a complementary and stronger filter than intensive soil tillage. Hence, OCA is hypothesized to allow the greatest potential for carabid beetle guilds to be expressed, followed by OA, then CA and CVA.

The present research goes beyond this preliminary study by covering the entire gradient between OCA and CVA, thus adding OA and CA, and detailing crop management practices to disentangle how each individual practice acts as an ecological filter influencing carabid communities. We conducted field sampling in 48 parcels in Belgium in 2021 and, provide a thorough analysis on the impacts of agricultural systems and practices on this crucial arthropod group. We hypothesize that different agricultural systems present different community compositions and diversity patterns because they involve different ecological filters. We expect alternative agricultural systems to support higher carabid diversity because reduced management-related filters likely increase the availability of ecological niches. More specifically, we expect that synthetic input use would act as a complementary and stronger ecological filter than intensive soil tillage. Consequently, we predict a diversity gradient from the most diverse to the least diverse carabid communities as follows: OCA > OA > CA > CVA. In contrast, we do not expect strong differences in carabid abundances between systems, as previous studies have reported inconsistent effects of farming practices on abundances, due to pitfall traps primarily reflecting activity-density rather than true abundances^23,24^ and some species being particularly adapted to conventional farming practices^12,25^.

## Material and methods

### Study site description

We conducted this study in Northen Wallonia, in Belgium (Figure 2). We included the ‘Condroz’ region but avoided the very contrasted (strong topography and landscape heterogeneity, cooler climate and shallow soils) environment of Southern Belgium (‘Ardennes’)^26^. The climate is temperate with a yearly average temperature of 11°C varying between an average of 3.7°C in January and an average of 18.4°C in July^27^. The sampling took place between March and July 2021, which showed temperatures below average values (average of 17,9°C vs 18,7°C usually) and relatively important rainfalls in July (average of 166,5 mm vs 76,9 mm usually). Agricultural systems in this region are predominantly conventional arable systems, characterized by the use of synthetic pesticides and regular soil ploughing, whereas organic farming and conservation agriculture remain in minority.

**Figure 2:**
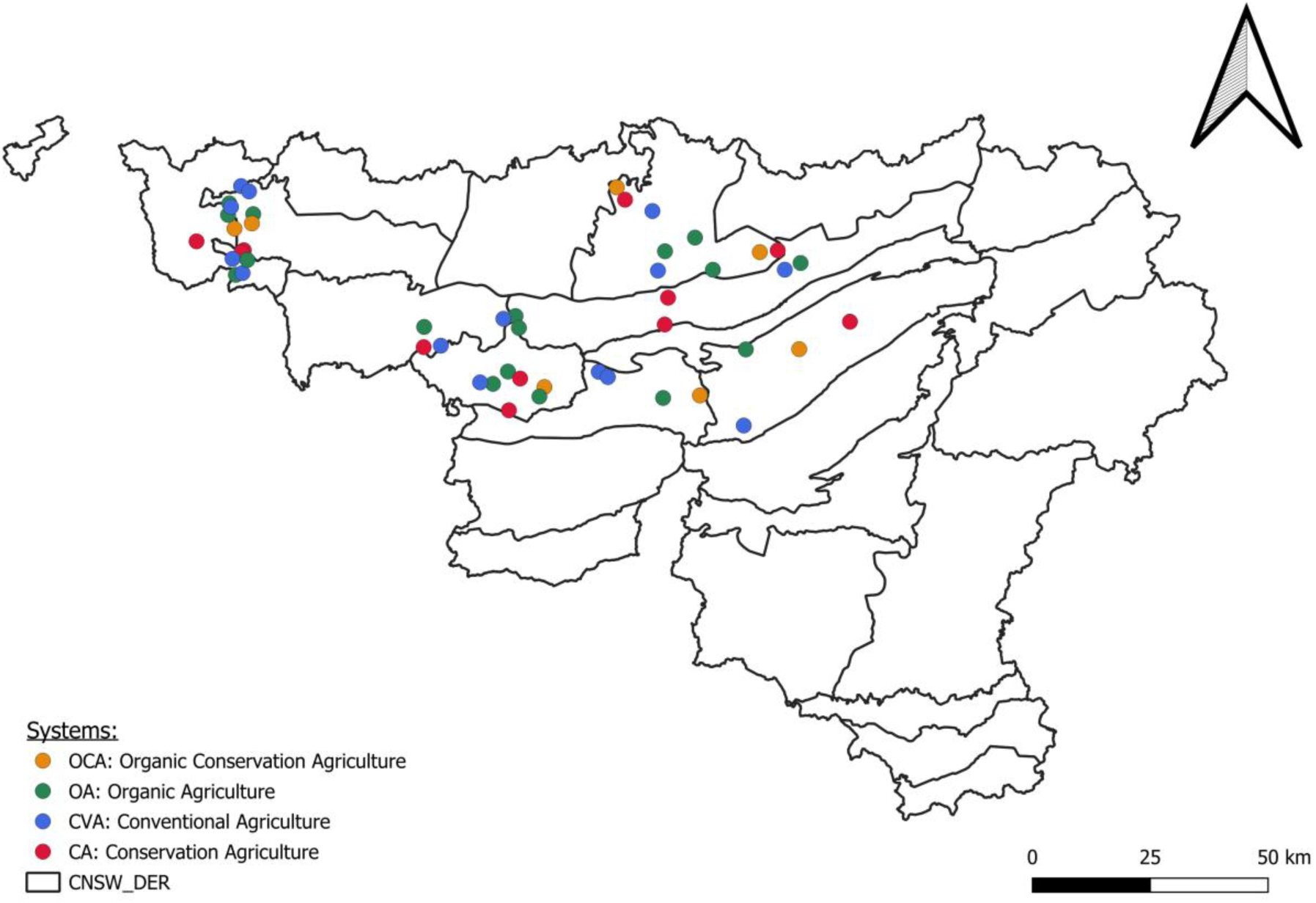
Sampling field locations across Wallonia region (Belgium). Agricultural systems are represented in distinct colors (see legend) and are evenly distributed across rural districts (i.e., CNSW_DER).

### Farm and parcel selection

We selected 48 farms dispersed over Northern Wallonia to ensure a homogeneous distribution of agricultural system categories across the landscape, each system category was represented evenly in each ‘rural district’, which are geographic units delineated based on eco-pedological parameters (rocks, landforms, soils and land use) (Figure 2).

All selected farms were commercial (i.e. non-experimental) cereal farms. Farms under OA (N = 17) were farms certified by the European Organic certification scheme, which prohibits the use of synthetic inputs (EU, 2007). Farms applying CA (N = 11) are farms relying on the three principles of CA (FAO, 2018a), i.e.: (i) the reduction of soil tillage intensity, (ii) the integration of cover crops in between cash crops and (iii) the diversification of crops within the rotation or within the parcels with intercropping. Farms in CVA (N = 14) do not comply with any organic certification scheme and do not apply the three CA principles. Farms in OCA (N = 6) are organically certified and apply the three principles of CA (Table 1).

**Table 1:** Classification of the four agricultural systems studied: OCA, OA, CA, CVA, and how each relates to the organic certification scheme and the application of the three CA principles. CA’s three principles: reduced tillage without soil inversion, the use of cover crops between cash crops, and crop diversification through rotation or intercropping.

|  | Organically certified | No organic certification |
| --- | --- | --- |
| Applying the three principles of CA | Organic conservation agriculture (OCA) | Conservation agriculture (CA) |
| Not necessarily applying the three principles of CA | Organic agriculture (OA) | Conventional agriculture (CVA) |

Within each selected farm, we selected one parcel of minimum 1 ha to avoid the border effect and minimum 1 km between them to avoid spatial correlation – this distance being conservative since carabids movements rarely exceed 10 m/day^28–30^. All selected parcels were growing winter cereals (avoiding winter barley, which has a distinct cropping calendar) at the time of sampling.

### Carabid sampling

We used two types of traps to sample carabids: pitfall traps and emergence traps. These two types of traps provide complementary information: pitfalls identify the density of activity, while emergence traps determine the hosting capacity of the parcel as an overwintering site as they capture overwintering pre-imaginal individuals^31,32^. Pitfall traps are traps made up of a container (dimensions: 400 mL, h=100 mm, top/inf diameter= 85/72 mm) buried at ground level to capture individuals running on the soil surface. A lid was placed approximately 20 cm above the container to prevent the pot from being flooded. Each trap was half filled with white vinegar, being a preservative solution, not attractive for the insects and without surface tension thus limiting the risk of escaping individuals. A 25 mm grid was placed above each trap to avoid the capture of micro-vertebrates. Emergence traps consist of tents hermetically sealing 0.36 m^2^ while collecting insects that emerge in that area with (i) one pitfall trap placed within the tent and (ii) one hanging jar collecting small flying carabids that do not usually fall into pitfall traps (Soil Emergence trap 96×26 mesh, Black, MegaView Science Co., Ltd., Taichung, Taiwan).

In each parcel, four pitfall traps and two emergence tents were placed 10 m apart in a row to minimize interference with mechanical interventions on the parcel. Each trap was at least 10 m from all edges of the plot. All traps were placed early-March and collected every two weeks until end of June 2021, which corresponds to 8 sampling events (i.e., T1-T8) for each parcel. All captured insects were stored in the refrigerator before sorting, after which collected individuals were stored in alcohol (96◦) for further identification with official identification keys^33,34^, using Fauna Europaea (https://fauna-eu.org) as a taxonomy standard.

### Practice description

Information about agricultural practices was collected through interviews with farmers in a standardized way. Information was gathered about practices applied before sowing of the cereal, and practices applied during the growing of the cereal to represent perturbations along gradients of soil mechanical disturbance and chemical input intensity. In total, interviews led to collecting data on 10 variables describing crop management (Table 2).

**Table 2:**
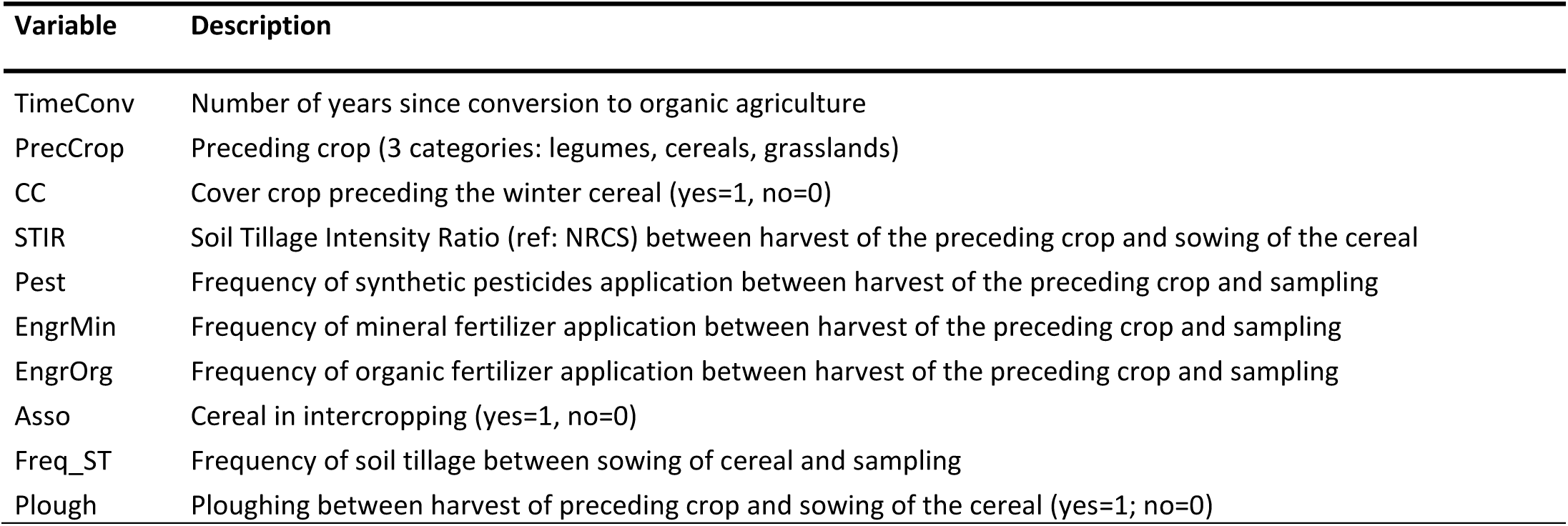
Variables used to describe agricultural practices derived from interviews with farmers.

### Statistical analyses

The data from the six traps were collated for each plot for the purposes of analysis. All analyses were computed in R version 4.5.1^35^.

Multivariate analyses were conducted to evaluate differences in species composition among across sampling sessions (T1-8) using Non-Metric Multidimensional scaling (NMDS) implemented in the R package *vegan*^36^. Prior to NMDS ordination, species abundance data were log-transformed (log(x+1)) to reduce the influence of highly dominant species while retaining zero values. NMDS was performed in three dimensions (k = 3), with a maximum of 100 random starts (trymax = 100). Ordination quality was assessed using the stress value below 0.20 which is considered acceptable representation of the dissimilarity structure. To test the environmental filtering hypothesis over the carabid community composition, the four agroecosystems were reclassified into two binary variables reflecting the system types: organic certification (1 = applied, 0 = absent) and conservation principles (1 = applied, 0 = absent) covering the four system categories as defined in Table 1. To control the effect of the sampling session on the carabid community (Figure 3A; Figure S2; Figure S3), we conducted a partial distance-based Redundancy Analysis (dbRDA) including agroecosystem classification (Table 1) as explanatory variables, with timing session a conditioning variable. After using *rankindex* function in vegan, we determined that Bray-Curtis index was the most appropriate dissimilarity measure for the community data. We reduced the relative differences between dissimilarities by applying a square-root transformation to the dissimilarity matrix. The statistical significance of the constrained model was assessed by permutation tests (ANOVA-like procedure) both by term and by axis, using 999 unrestricted permutations.

**Figure 3:**
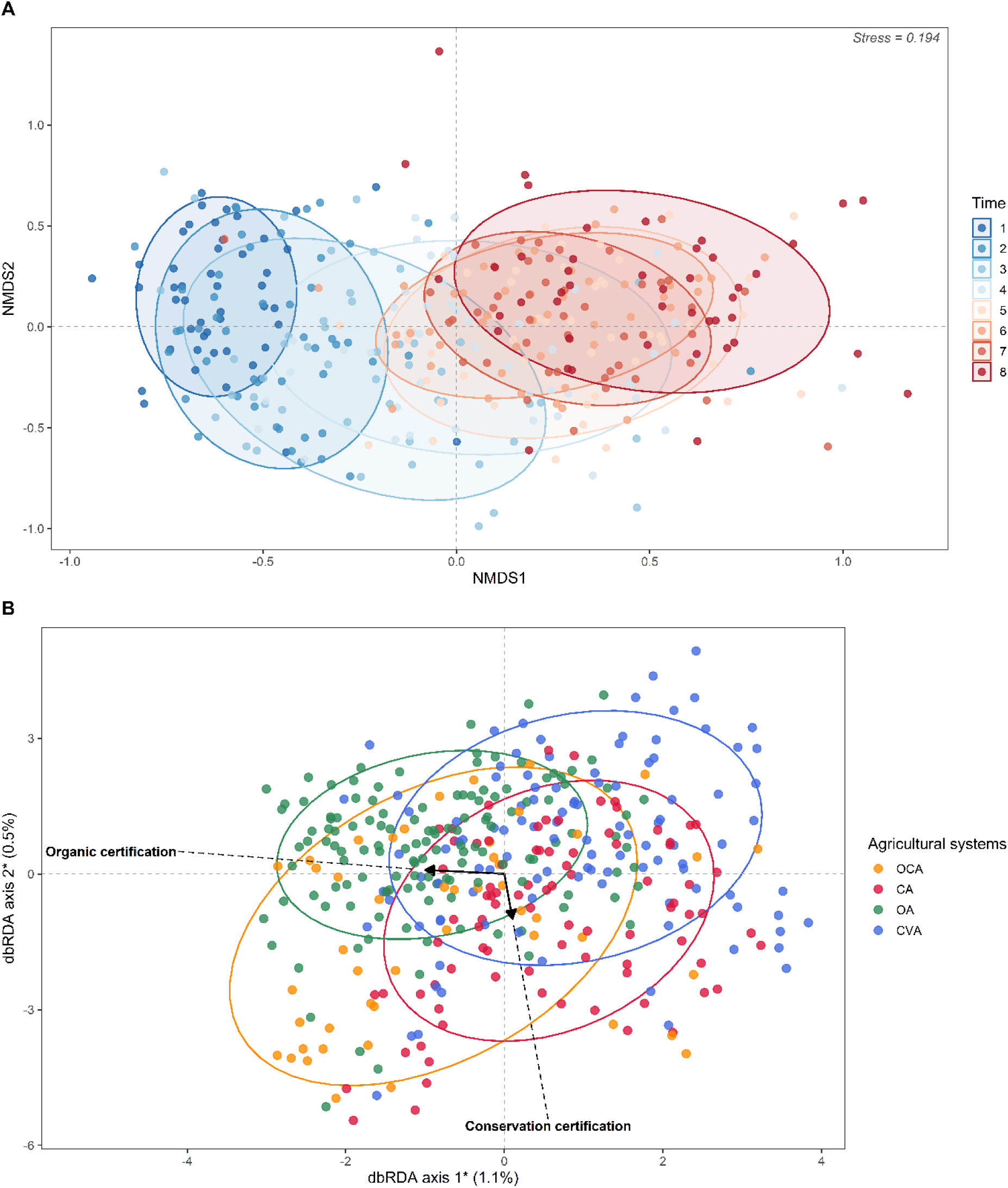
(A) Non-metric multidimensional scaling (NMDS) ordination of carabid communities based on Bray-Curtis dissimilarities computed on log-transformed abundances (stress = 0.194, k = 3) with sampling sessions (T1-8) colored along a color gradient. Each point represents a sampled plot at a given sampling session. (B) Partial distance-based Redundancy Analysis (dbRDA). Figure dots are colored by agricultural system depending on their binary definition following Table 1: organic-conservation (OCA, yellow), organic (OA, green), conservation (CA, red) and conventional agriculture (CVA, blue). Axes represent the constrained canonical axes, with values indicating the proportion of total variance explained by each axis. Ellipses show 80% confidence around the centroid of each sampling session (A) and each agricultural system (B). * corresponds to significant axis.

Then, permutational multivariate analysis of variance (PERMANOVA) was subsequently applied to identify which agricultural systems (i.e., OCA-OA-CA-CVA) supported significantly distinct carabid communities using the *adonis2* function of *vegan* R package^36^. To maintain consistency with the partial dbRDA framework and control for phenological variation, the effect of sampling session was first removed by fitting a dbRDA constrained solely by *Time* using the Bray-Curtis dissimilarity index with square root transformation of the dissimilarity matrix. Pairwise comparisons between all six combinations of agricultural systems levels were subsequently performed using the *pairwise.adonis2* function of the R package *pairwiseAdonis*^37^, with p-values adjusted for multiple comparisons using the Benjamini-Hochberg false discovery rate correction. All permutation tests were conducted with 999 unrestricted permutations. To quantify the relative and independent contributions of organic certification, conservation status, and sampling sessions to the variation in carabid beetle community composition, a variation partitioning analysis was performed using the *varpart* function of the *vegan* package^36^.

Finally, species accumulation curves were computed using the function *specaccum* of R package *vegan*^36^ to estimate the total carabid species richness within each agricultural system from the non-transformed community matrix of carabids.

From here, community data were aggregated across sampling sessions, to retain a single sample per plot and avoid farm pseudo-replication. Alpha diversity metrics, including carabid richness, carabid abundance, Hill-Shannon’s and Chao bias-corrected indexes (i.e., the estimated carabid species richness), were calculated for each sampled farm. The Hill-Shannon’s index, which represents the effective number of species based on Shannon’s entropy, was estimated using *iNEXT* R package ^38^. This index minimizes the weight of dominant species. Chao bias-corrected^39^ was estimated using “*estimateR*” function in the *vegan* package^36^ to estimate the “true” number of species in the community based on the community matrix of carabid beetles. To compare the generated alpha diversity metrics across agricultural systems, Linear Models (LMs) and Generalized Linear Models (GLMs) were performed using MASS R package^40^. All models included agricultural system as a fixed effect. Observed carabid richness was analyzed using a Poisson error distribution. For carabid abundance, a negative binomial error distribution was used due to overdispersion. Chao bias-corrected richness and the Hill–Shannon diversity index were analyzed using linear regression models. Post hoc pairwise comparisons among agricultural systems were performed using Tukey-adjusted contrasts implemented via the *glht* function in the *multcomp* R package^41^.

To evaluate the influence of specific agricultural practices on species composition, another dbRDA was applied using Gower dissimilarity index. We checked the collinearity between our quantitative variables (thus removing PrecCrop). Multicollinearity among agricultural predictors was assessed using variance inflation factors (VIF) via the *vif.cca* function. Variables with VIF values exceeding our threshold (e.g., > 3) were removed^42^. The overall significance of the constrained ordination model was tested using PERMANOVA with 999 permutations. Significance was then assessed separately for each canonical axis and for each agricultural predictor (999 permutations).

Because a single species dominated the dataset (44%), *Pterostichus cupreus* (Linnaeus, 1758), we repeated all multivariate analysis after excluding it, to check that the detected effects of agroecosystems and agricultural practices were not driven by this dominant species alone.

To evaluate the influence of specific agricultural practices on alpha diversity measure (i.e., carabid observed richness, estimated richness, abundance, and Hill-Shannon Index), we used the same model construction of the agroecosystem influence over alpha diversity indexes. The full model for each diversity metric included all agricultural practices as predictors from Table 2. Then, we conducted multi-model inference based on AICc (Akaike Information Criterion corrected for small sample sizes) to evaluate support for candidate models. We performed model dredging using the *dredge* function from the *MuMIn* package^43^, fitting all possible subsets of fixed effects from the global LM or GLM. We ranked models by AICc and retained the best model. We retained the best selected model (top 1) for each alpha diversity measure fitting the same structure as for the previous modelling analysis. All model assumptions were verified using the *DHARMa* R package^44^.

## Code and data availability

The underlying raw data and code for this study is available in Figshare repository and can be accessed via this link https://doi.org/10.6084/m9.figshare.34027950.

## Results

We recorded a total of 33,943 carabid specimens belonging to 27 genera and 83 species from 48 agricultural parcels across the four agricultural systems. Nearly half of our records (N = 14,796) were dominated by *P. cupreus* (Figure S1).

### Phenology, organic certification and soil conservation principles as ecological drivers of carabid composition among agricultural systems

NMDS revealed a temporal gradient along the first axis from earlier sampling sessions (T1) on the left and later ones (T8) on the right (Figure 3A), indicating an influence of seasonality on carabid composition. From T1 to T4 (March until end of April; blue colors), species composition varies between sampling, while from T5 (May) until T8 (June), species compositions overlap (red colors), indicating greater similarity among the last four sampling sessions. Our variation partitioning analysis (adjusted R² of residuals = 93.4% ) showed that sampling session (adjusted R² = 5.6%), organic status (adjusted R² = 0.8% ) and conservation status (adjusted R² = 0.2%) can be considered as independent variables given that shared variance accounted for nearly 0% (Figure S2).

The partial dbRDA, controlling for sampling sessions (5.84% of total variance conditioned), revealed that organic agriculture and conservation agriculture jointly explained 1.51% of the constrained variation in carabid community composition (Figure 3B). The overall constrained model was significant (df = 2; F = 3.06; p < 0.001), with both organic agriculture (df = 1; F = 4.278; p < 0.001) and conservation agriculture (df = 1; F = 1.841; p = 0.002) independently contributing to community differentiation. Organic agriculture exerted an effect nearly 2.32 times greater than conservation agriculture based on both F-values and sums of squares, highlighting the predominant role of chemical inputs over physical soil disturbance in structuring carabid assemblages.

PERMANOVA showed a significant difference across systems (df = 3; R^2^ = 0.024; F = 3.12; p-value = 0.02). Paired comparisons showed significant contrasts between OCA with CA and CVA, and between OA and CVA (Table 3). In addition, the magnitude of the effects followed a consistent pattern. The strongest differences (highest pseudo-F and lowest adjusted p-values) were systematically found between organic (OA, OCA) and conventional (CVA) systems, the contrast between OCA and CVA being the most pronounced. Conversely, the two comparisons between systems differing only in the application of conservation practices — OA versus OCA (both organic) and CA versus CVA (both non-organic) — showed weak differentiation suggesting that the organic status primarily drives community composition. OA and CA did not show significant differentiations indicating that these two single-filter systems supported largely indistinguishable assemblages despite differing in their filter type.

**Table 3:** Outcomes of statistical test of variance analyses (PERMANOVA) between distance matrices of each agricultural system (degree of freedom of the model = 1). Bold characters mean that the paired comparison was significant after Benjamini-Hochberg correction (adjusted p-value).

| Paired comparison | F | R<sup>2</sup> | Adjusted p-value |
| --- | --- | --- | --- |
| CA vs CVA | 0.640 | 0.003 | 0.502 |
| CA vs OA | 1.530 | 0.007 | 0.222 |
| **CA vs OCA** | **4.074** | **0.030** | **0.035** |
| **CVA vs OA** | **4.486** | **0.018** | **0.018** |
| **CVA vs OCA** | **7.222** | **0.045** | **0.007** |
| OA vs OCA | 1.60 | 0.009 | 0.195 |

The exclusion of *P. cupreus* from the dataset did not significantly alter the outcomes of the explained variance of partial db-RDA but the subsequent pairwise PERMANOVA showed an increased magnitude of most paired comparisons and an uncovering of two more significant paired-comparisons, indicating that this dominant taxon may drive some patterns of differentiation among system types (Figure S4; Table S1).

### Agricultural systems influence species diversity

Accumulation curves show that OCA and OA reached higher species richness than CA and CVA, indicating that organic farming is associated with higher diversity levels (Figure 4).

**Figure 4:**
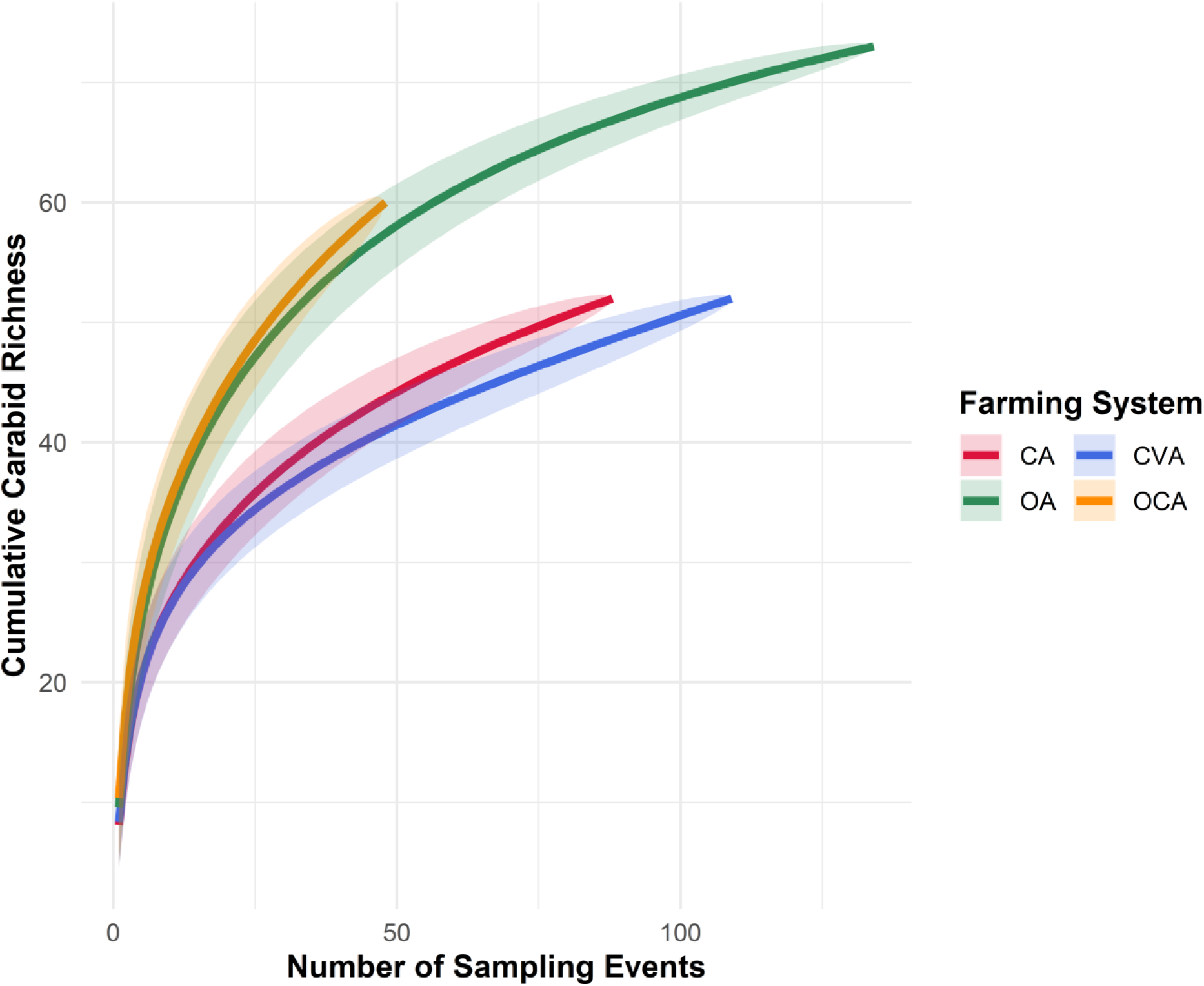
Accumulation curves per agricultural system: Conservation Agriculture (CA) in red, Conventional Agriculture (CVA) in blue, Organic Agriculture (OA) in green and Organic Conservation Agriculture (OCA) in yellow. Shadows correspond to confidence intervals of each accumulation curve.

We observed significant differences in alpha diversity metrics among agricultural systems (Figure 5). Both observed and estimated carabid richness were significantly higher in OCA than in CVA (observed richness: estimate = 0.257, z-value = 2.719, *p-value* = 0.033; estimated richness: estimate = 15.267, z-value = 2.662, *p-value* = 0.05) and than in CA (observed richness: estimate = 0.256, z-value = 2.590, *p-value* = 0.047; estimated richness: estimate = 17.277, z-value = 2.896, *p-value* = 0.028). Additionally, observed richness was significantly higher in OA than in CVA (estimate = 0.198, z-ratio = 2.717, *p-value* = 0.033). In contrast, carabid abundance and dominance–rarity patterns (Hill–Shannon index) did not differ significantly among agricultural systems.

**Figure 5:**
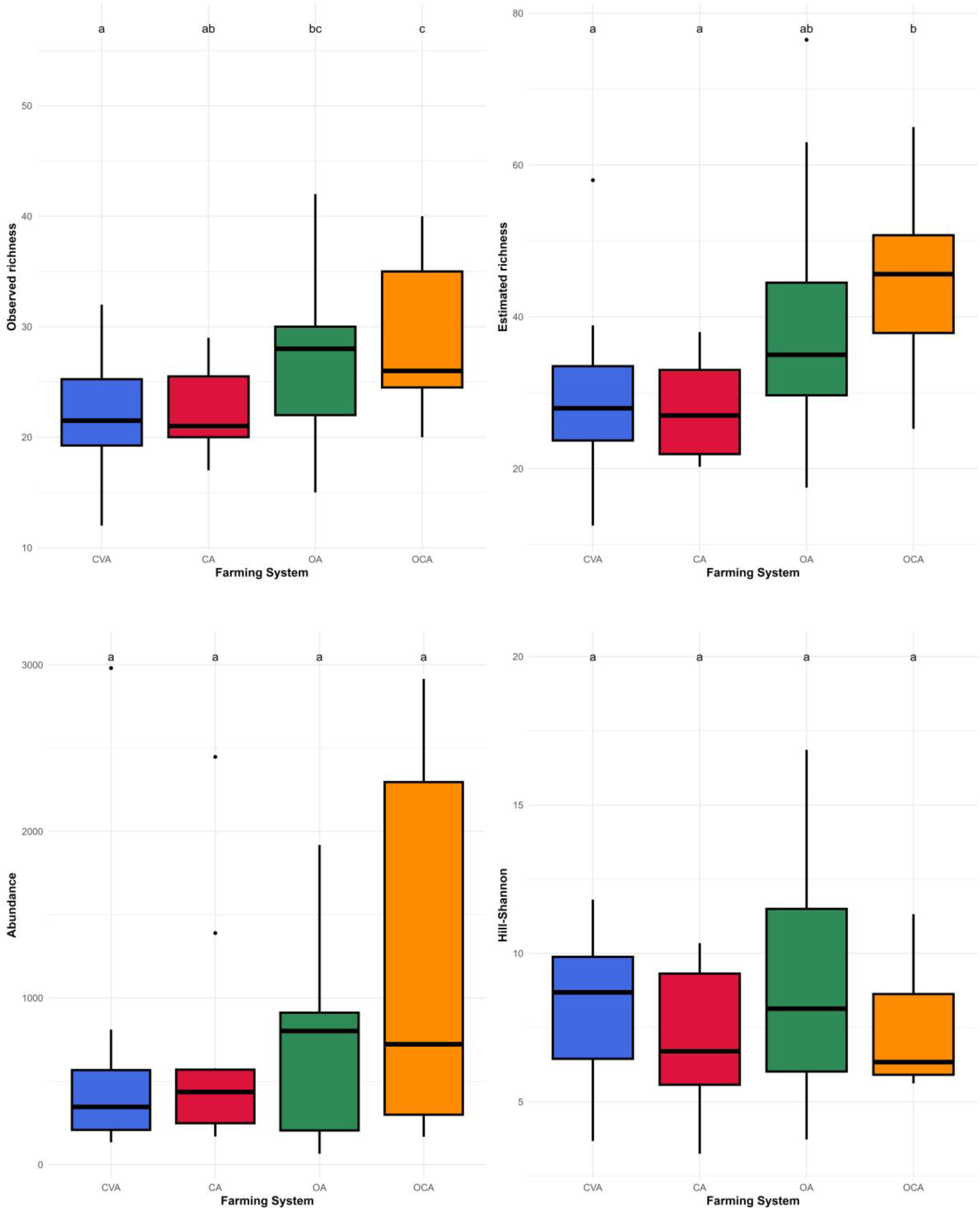
Agricultural systems comparisons among Conservation Agriculture (CA) in red, Conventional Agriculture (CVA) in blue, Organic Agriculture (OA) in green and Organic Conservation Agriculture (OCA) in yellow according to the observed carabid richness, estimated carabid richness, carabid abundance, and Hill-Shannon’s index.

### Agricultural practices explain differences across communities

Analyzing communities at the practice level allows for a more detailed understanding of the specific drivers involved. Among the quantitative variables kept for analysis, the frequency of mineral fertilizer was removed from our analysis due to multicollinearity (VIF > 3). In particular, dbRDA analysis revealed that the eight agricultural practices significantly explained 22.5% of the constrained variance on carabid community composition (df = 8, F = 1.416, *p-value* = 0.008; Figure 6). The first constrained axis contributed 40.59% to this explained variance, corresponding to 9.1% of the total variance (i.e., constrained and unconstrained), and was statistically significant (df = 1, F = 4.597, *p-val*ue = 0.005). Among the eight predictor variables, two significantly influenced carabid community dissimilarity: the frequence of synthetic pesticide application (df = 1, F = 2.810, *p-value* = 0.001) and time since conversion to organic agriculture (df = 1, F = 2.656, *p-value* = 0.009). The exclusion of *P. cupreus* did not alter the outcomes of the dbRDA except by slightly diminishing the constrained variance (20.53%) of the agricultural practices’ variables and canonical axis (Figure S5) indicating that the detected effects were robust and not driven by this dominant species alone.

**Figure 6:**
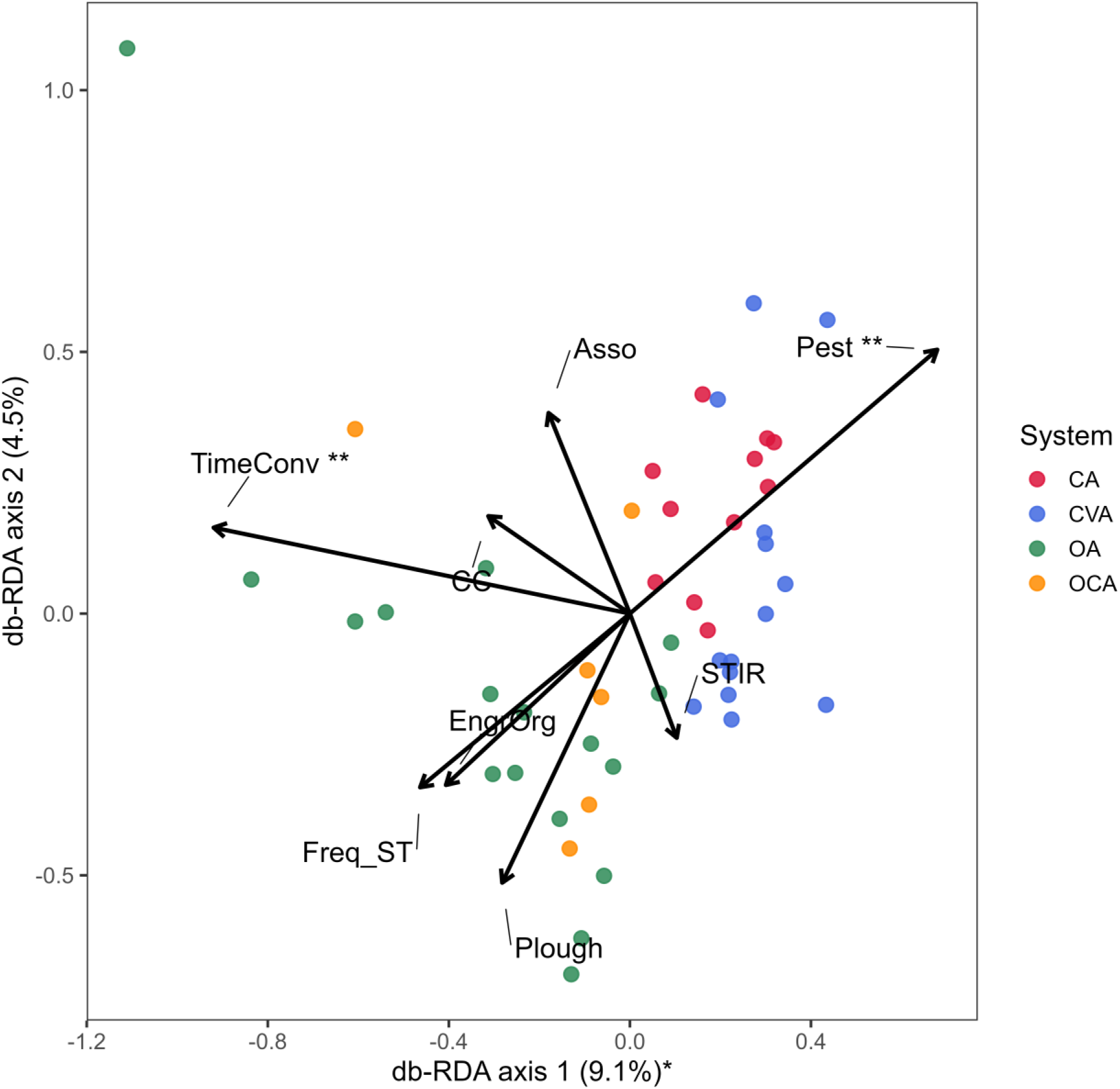
Distance-based Redundancy Analysis (dbRDA) ordination of carabid communities across agricultural fields. Colors correspond to agricultural systems: Conservation Agriculture (CA, red), Conventional Agriculture (CVA, blue), Organic Agriculture (OA, green), and Organic Conservation Agriculture (OCA, yellow). Axes represent the constrained canonical axes, with values indicating the proportion of total variance explained by each axis. Asterisks denote statistically significant: * for *p-value* < 0.05 and ** for *p-value* < 0.01. Black arrows represent quantitative predictor variables of agricultural practices, with arrow length and direction indicating the strength and sign of their influence on community composition.

We found that time since conversion to OA, the presence of cover crops, and the frequency of organic fertilizer application had positive effects on observed carabid richness (Table 4). The frequency of mineral fertilizer application had a negative effect on estimated carabid richness. The frequency of synthetic pesticide application and the occurrence of ploughing before sowing had significant negative effects on carabid abundance. Hill Shannon index was not influenced by any of the predictors, suggesting that agricultural practices have no influence on the dominance-rarity pattern of the carabid community.

**Table 4:** Models tested for each alpha diversity indices, corresponding AIC values and p-values for each model predictor. The definitions of each selected predictor are given in. **Table 2**.

| Model Response | Weight of Best model - AICc.w | Selected Predictors | Estimate ± SE | $p\text{-value}$ | $R^2$ |
| --- | --- | --- | --- | --- | --- |
| Observed carabid richness | 0.24 | TimeConv | 0.014±0.005 | **< 0.001** | 0.075 |
| | | CC | 0.128±0.059 | **0.03** | |
| | | EngrOrg | 0.084±0.042 | **0.04** | |
| Estimated carabid richness | 0.42 | EngrMin | -3.757±1.251 | **0.004** | 0.16 |
| Carabid abundance | 0.26 | Pest | -0.16±0.047 | **< 0.001** | 0.013 |
| | | Plough | -0.473±0.239 | **0.048** | |
| Hill-Shannon Index | 0.44 | ~ 1 | 7.925±0.421 | **< 0.001** | NA |

## Discussion

While most research addressing the impact of agriculture on biodiversity has so far focused on the system type level, often comparing one alternative system (mainly OA) to CVA, this research is pioneer by comparing multiple alternative systems and by providing a detailed understanding of how agricultural practices associated with these different agricultural systems influence carabid communities.

### Carabid community composition is primarily driven by seasonality from March until April

Our results show that species composition varied across early sampling sessions, while it progressively converged later in the season. This result is consistent with previous work that highlighted the strong influence of seasons on carabid communities^45^, and their interaction with agricultural management^46^. It likely reflects the phenology of the community with early spring samples capturing different overwintering adults species through time reflecting diverging species thermal requirements and reproduction cycles^47^. As the season progresses, the assemblage becomes increasingly dominated by a few dominant species (such as *P. cupreus*, a spring breeder) which homogenizes samples. Yet, it is to be noted that this particular timing observed may be specific to 2021 which was cooler than averages which may have influenced emergence and activity peaks.

### Organic agriculture shapes carabid composition two times more than conservation agriculture, and supports higher carabid richness

Contrary to our prediction of four discrete significantly ordered levels (OCA > OA > CA > CVA), the agroecosystems could not be individually separated and the gradient was mainly supported at its extremes with OCA showing the most differentiated communities and richness levels compared to CVA. Community composition and richness levels therefore behaved as a continuous gradients rather than as a set of sharply delimited categories. The magnitude of the effect was mainly driven by the organic status which explained community structure 2.32 times more than the soil conservation status. Conversely, adding the conservation-agriculture principles (to OA, yielding OCA) or applying them alone (CA vs CVA) produced no significant gain in carabid richness or composition.

The strong differentiation of OCA with regards to both CVA and CA in terms of community composition and richness is likely driven by the absence of two major filters influencing carabid communities: synthetic inputs and intensive soil tillage. This result was expected as the absence or decrease of ecological filters likely increases ecological niches, therefore allowing diversity to be expressed. While OCA systems have been little studied, our results corroborate previous studies: Rivers *et al.* (2017, 2018)^48,49^ studied CA transitioning towards OCA and showed that the activity-density, species richness and evenness of carabid populations increased throughout the transition, thus showing strong positive potential of OCA. More recently, Boeraeve et al. (2022)^21^ highlighted higher alpha, beta and functional diversity carabids in OCA as compared to CVA. While evidence remains limited, these results so far appear consistent. Yet, it should be acknowledged that combining both approaches is agronomically complex^18^ as the absence of ploughing, mineral fertilizers and herbicides usually leads to reduced mineralization^50^ and increased weed competition^51^.

The overall tendency of the organic status to host higher species richness than CVA, has been previously documented for biodiversity overall^52,53^ and carabids in particular^1,54,55^. This illustrates the main ecological filter role played by pesticides. While a small share of studies nuance this general trend (e.g. the meta-analysis of Bengtsson et al. (2005)^1^ found 16% studies to have a negative effect of OA on species diversity), which can be explained by methodological bias^24^ and the numerous interacting factors^56–58^, this positive effect of OA on biodiversity, and specifically carabids, seems nowadays to reach a scientific consensus.

The absence of response of CA with regards to CVA in terms of carabid composition (i.e., including *P. cupreus* population), abundance and diversity contrasts with the large body of literature that reports a positive outcome of conservation practices on carabid abundance and diversity^6,14,22,59–61^. Yet, this literature is often restricted to the first pillar or the three conservation principles, i.e. soil tillage reduction. In terms of reduction of soil disturbances, it is indeed expected to have a positive impacts on carabids, for which part of the life cycle takes place underground^60^. Other, less direct explanations include increased organic matter or seeds on the soil surface increasing food and niches availabilities^13^. However, the lack of a clearcut definition and the absence of commercial specifications means that CA can be associated with a wide range of management practices^20,59^ which was reflected in our practice data with strong heterogeneity in tillage intensity and cover cropping for instance. This may explain the high variability of carabid responses to CA and further supports practice-level analyses as carried out in this research.

Interestingly, although agricultural systems influenced species richness and community composition, they did not significantly affect abundances and Hill–Shannon diversity index indicative of the overall dominance structure of communities. One possible explanation is the strong dominance of a few widespread generalist species across all systems, particularly *P. cupreus*, which represented nearly half of all captured individuals.

### From systems to practices: pesticide use and ploughing impact abundances while time since organic certification, cover crops and organic amendment drive richness

Looking at the practices levels brings more detailed information on which precise practice drivers act positively and negatively on the distinct biodiversity indicators and how practices act in tradeoffs or synergies. Interestingly, the two practices that significantly reduced carabid abundance — synthetic pesticides and ploughing — tend to substitute for one another across systems: removing the pesticide filter under organic management comes often at the cost of maintaining, or even intensifying, ploughing, because weeds can no longer be controlled chemically. In fact, in our farm sample, OCA farms were ploughed before sowing as often as CVA ones. This trade-off may thus help explain why we detected no difference in abundance among agricultural systems, even though abundance was clearly sensitive to these individual practices. Richness, in contrast, was driven by practices associated with organic management that are not caught in this trade-off: time since conversion, cover crops and organic amendments. This may explain why the benefits of OA and OCA were expressed more clearly on richness than on abundance.

The use of use of organic amendment^62^ and the absence of synthetic pesticides^63^ are both acknowledge in the literature as influencing distinct carabid communities. Our study confirmed that organic amendment had a positive effect on richness and the quantity of pesticide on abundances. The quantity of pesticides^63^ and their toxicity^64^ are now widely acknowledged to be detrimental to carabids and other non-target soil organisms such as earthworms^65^ or collembola^66^.

Moreover, our study confirms that time since conversion has a significant effect on carabid communities. Indeed, the longer the farm had converted to organic agriculture, the higher its richness, and it also significantly structured community composition. This is consistent with previous work that showed positive influence of OA over time^67^ possibly reflecting the gradual recovery and stabilization of carabid communities under long-term organic management.

### Perspectives

It is worth noting that agricultural systems and measured practices together explained only a modest share of the compositional variation, leaving the large majority attributable to factors we did not quantify: local climatic and microclimatic conditions (particularly relevant given the atypically cool 2021 season), local soil physico-chemical properties (texture, moisture, organic-matter content, pH) that condition the development of soil-dwelling pre-imaginal stages, and the surrounding landscape context (field margins, semi-natural elements, crop heterogeneity), a well-documented determinant of carabid communities. Because organic and conservation farms are not randomly distributed across the landscape or across soil types, some of these unmeasured variables may also act as confounders rather than merely as additional noise. Integrating such climatic, edaphic and landscape data represents a promising avenue for future research aiming to disentangle the respective contributions of farming practices and their environmental setting.

Our results also show a high variability of practices among farming systems encouraging future research to go beyond the system level. Sampled CA parcels were significantly less ploughed before sowing than the three other systems, but interestingly OCA systems, supposedly applying the conservation principles of CA, were ploughing as much as CVA and OA. Another example lies in the cover crop practice, being part of the three defining pillars of CA^11^, yet not more implemented in CA or OCA systems than other systems. Since our results show that cover crops have a positive effect on observed carabid richness, strengthening this pillar within CA systems seems like a promising avenue. The impact of cover crops on carabids has been little addressed in the literature, but a recent systematic review corroborates our results although it highlights that this beneficial effect varies across cover crop species, termination time and methods, and cover crop management^68^.

While it is of interest for research to compare agricultural systems and understand their relative impacts on carabid communities, it is also critical to acknowledge that there is no one-size fits all solution across taxonomic groups. Indeed, a 14 year-old study focusing on soil organisms in OA, CA and CVA showed a higher overall improvement in CA than in OA^2^. It is thus likely that the relative role of practice filters vary across organisms and that the impact of agricultural systems depends upon the targeted organism’s traits and life cycles characteristics. Carabids, as mostly epigeal, surface-active predators, appear to benefit primarily from the removal of the pesticide filter under organic management, whereas soil-dwelling organisms may respond more strongly to reduced soil disturbance under conservation agriculture, as suggested by Henneron *et al.* (2015) ^2^. This argues in favor of OCA systems which combine both filtering removals and may therefore benefit a broader range groups. However, if such integrative and systemic approach is appealing in terms of environmental sustainability, very few farmers endorse the challenge as it offers fewer levers to correct problems and requires anticipating them through system design rather than curative interventions, and little commercial valorization at the food chain level^69^. OCA still requires further research to support higher yield stability and food chain adaptation to remunerate risks undertaken by OCA farmers. Yet, research on alternative tillage methods for organic agriculture remains limited as compared to research directed at conventional agriculture^70^. Future policy and research efforts should therefore aim to foster synergies among agricultural systems.

## Conclusions

In order to better understand the drivers of biodiversity in agricultural systems, there is a need to differentiate among the wide variety of agricultural systems and practices they entail. In this study, we contribute to filling this gap by comparing the effect of four major agricultural systems on carabid communities and by exploring the role of 10 practices to understand these effects. We show that the that carabid communities are structured first by seasonality and, secondarily, by organic status rather than by tillage regime. Observed and estimated richness were higher under organic management, with organic-conservation agriculture (OCA) offering the greatest potential for carabid richness, while abundance and the dominance structure of communities (Hill–Shannon index) did not differ among systems. Moving from systems to practices clarified these patterns: carabid abundance was reduced by synthetic pesticides and ploughing, whereas richness was promoted by time since conversion to organic farming, cover crops and organic amendments. The negative effects of synthetic pesticides and ploughing likely tended to offset one another across systems, probably explaining why no difference in abundance emerged at the system level.

These results position pesticide use, time since conversion, cover crops, organic amendment and ploughing as the main levers acting on carabid communities and highlight the value of analyzing detailed practices rather than farming systems alone, given the strong heterogeneity of practices we observed within each system category. Our inferences, however, concern the taxonomic structure of assemblages (i.e. richness, abundance and composition) and not the pest-control function they may support, which we did not measure; trait-based approaches are a necessary next step before recommendations can be framed in terms of ecosystem services. Further research on alternative tillage methods for organic agriculture must expand to encourage the design of agricultural systems reconciling food production and all groups of biodiversity.

## Supporting information

https://doi.org/10.6084/m9.figshare.34027950

## Acknowledgments

We are grateful to all the farmers who participated in this study, for granting us access to their fields and for sharing detailed information on their management practices. This work would not have been possible without their collaboration. We thank XXXX for funding XX’s postdoctoral grant, and XX for his help in verifying the identification of specific carabid specimens. We also thank the anonymous reviewers for their constructive comments on the manuscript.

