## Supplementary material for "Carabid beetle communities along a gradient of agroecological systems: disentangling the roles of pesticides and soil tillage": https://doi.org/10.6084/m9.figshare.34027950

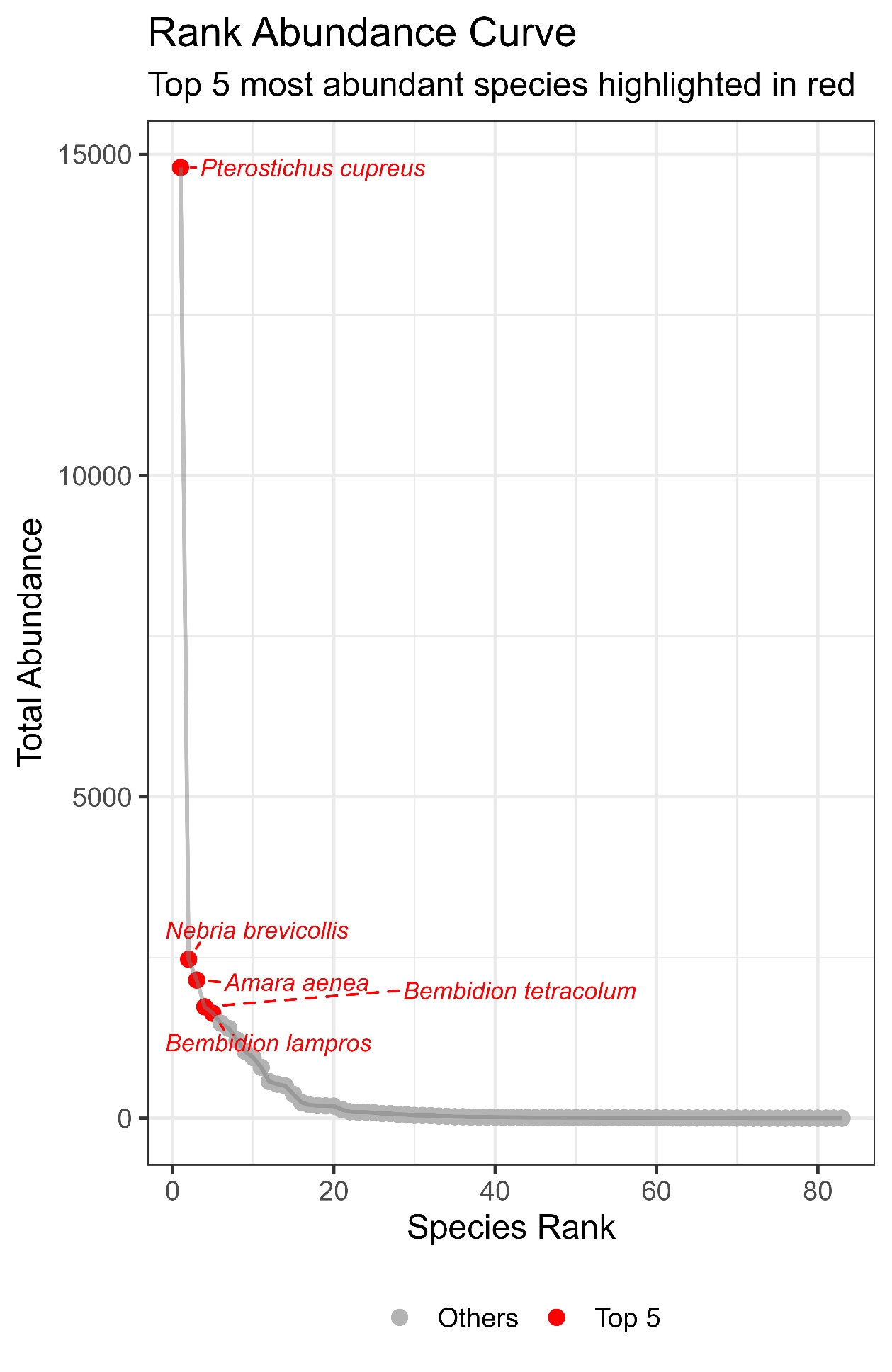

Figure S1 : Abundance rank curve showing total abundance of carabid species. The top 5 sampled species are highlighted in red.

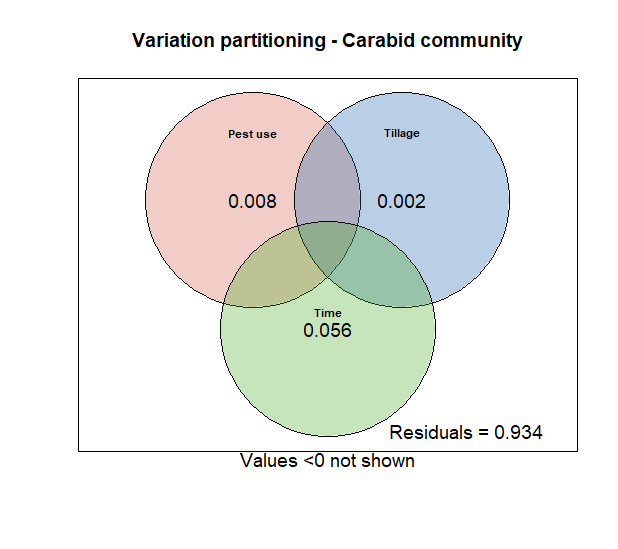

Figure S2 : Variation partitioning Venn diagram. The explained variance is within each circle for sampling session (green), pesticide use (red) and soil tillage (blue).

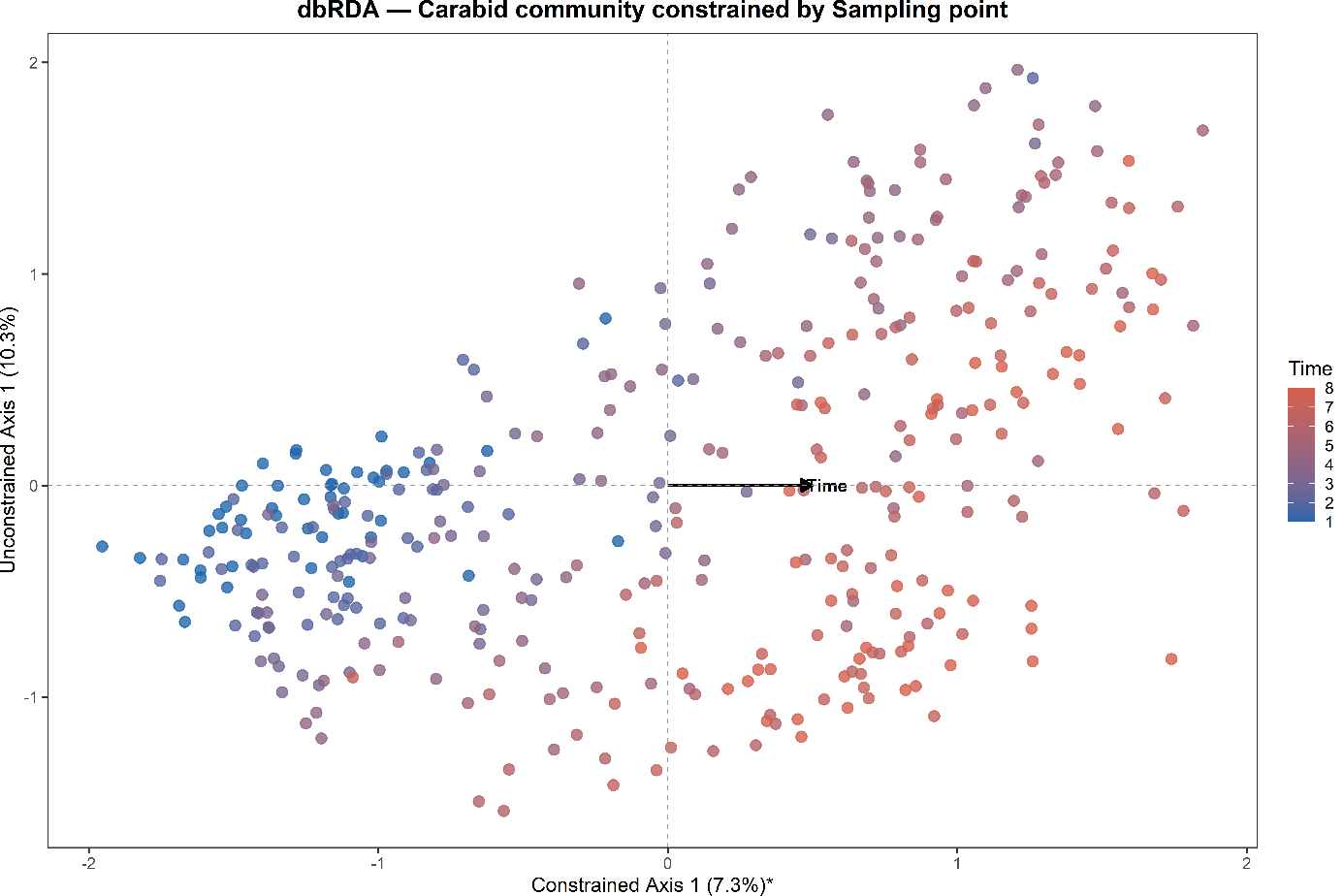

Figure S3. Distance-based Redundancy Analysis (dbRDA) only constrained by sampling points. Figure dots are colored by sampling session (T1-T8) along blue-red color gradient. First axis corresponds to the first constrained dimension and the second axis corresponds to the first unconstrained dimension with values indicating the proportion of total variance explained by each axis. * corresponds to significant axis.

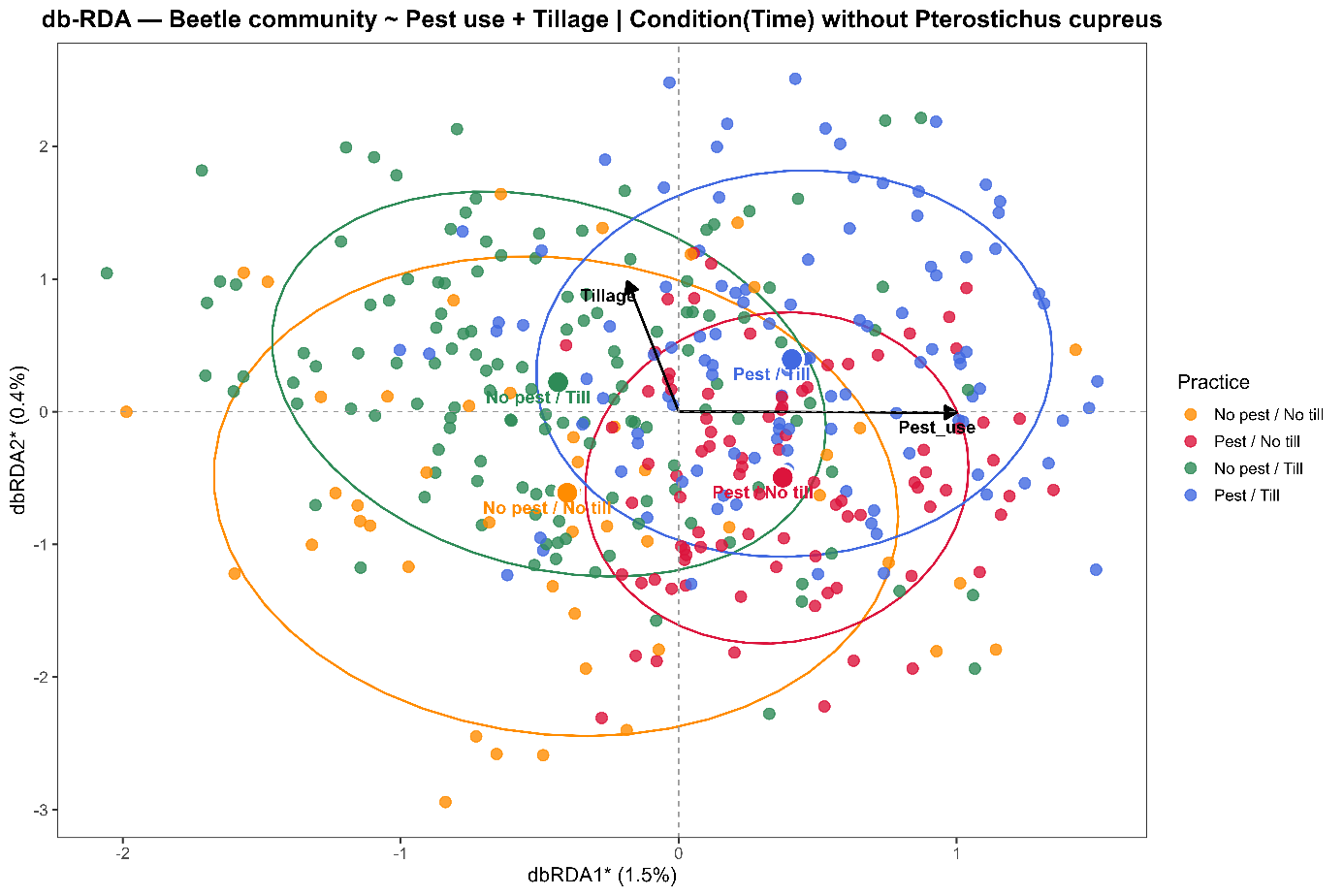

Figure S4.  Partial distance-based Redundancy Analysis (db-RDA) after removing *Pterostichus cupreus* (Linnaeus, 1758). Figure dots are colored by agricultural system depending on their binary definition following Table 1: organic-conservation (OCA, yellow), organic (OA, green), conservation (CA, red) and conventional agriculture (CVA, blue). Axes represent the constrained canonical axes, with values indicating the proportion of total variance explained by each axis.  Ellipses show 80% confidence around the centroid of each agricultural system. * corresponds to significant axis.

Table S1. Outcomes of statistical test of variance analyses (PERMANOVA) between distance matrices of each agricultural system (degree of freedom of the model = 1) after removing *Pterostichus cupreus* (Linnaeus, 1758). Bold characters mean that the paired comparison was significant after Benjamini-Hochberg correction (adjusted p-value).

| Paired comparison | F | R² | Adjusted p-value |
| --- | --- | --- | --- |
| **CA vs CVA** | **4.574** | **0.023** | **0.01**** |
| **CA vs OA** | **2.968** | **0.013** | **0.044*** |
| CA vs OCA | 2.231 | 0.016 | 0.012 |
| **CVA vs OA** | **4.784** | **0.019** | **0.005**** |
| **CVA vs OCA** | **4.579** | **0.029** | **0.007**** |
| OA vs OCA | 0.266 | 0.001 | 0.764 |

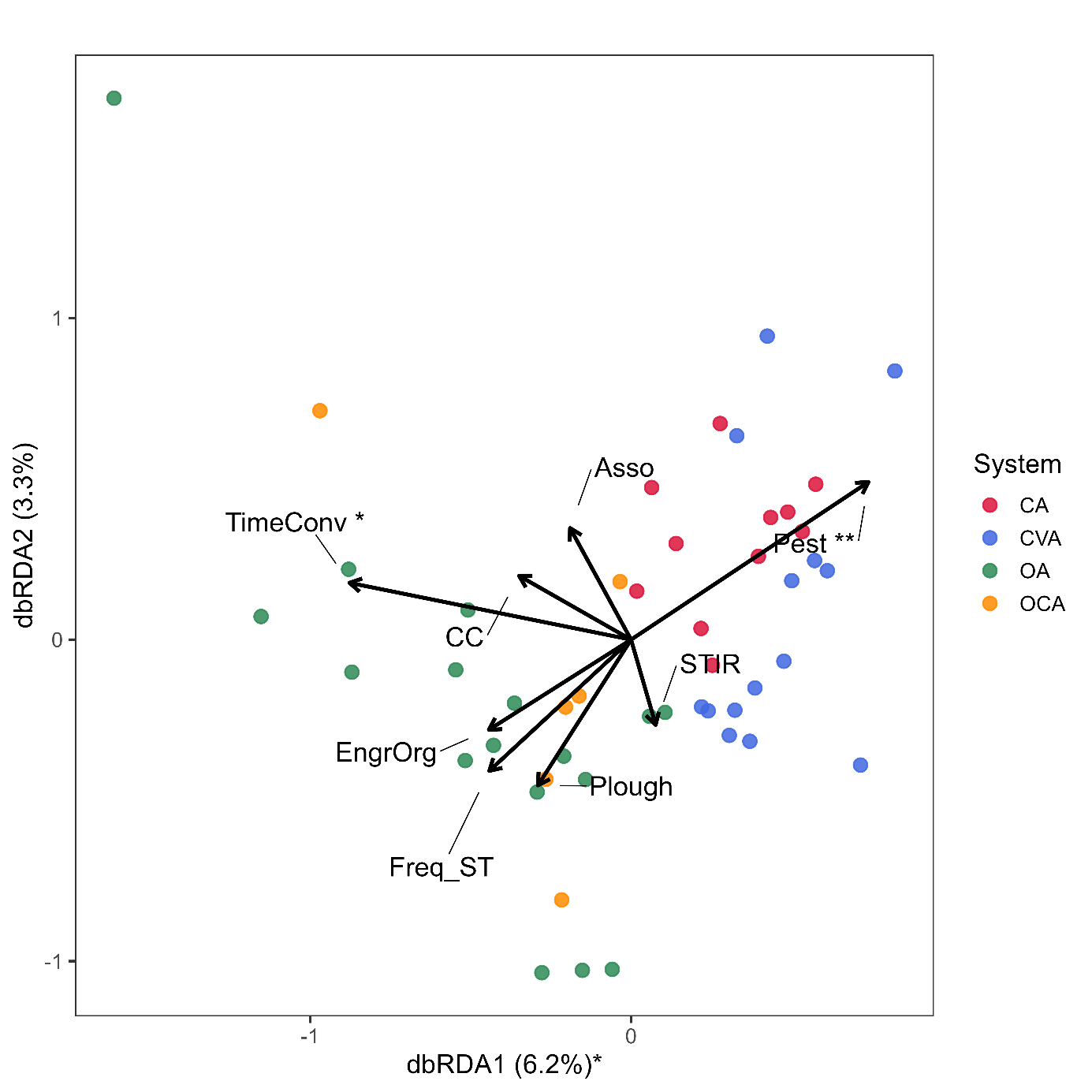

Figure S5. Distance-based Redundancy Analysis (db-RDA) ordination of carabid communities across agricultural fields after removing *Pterostichus cupreus* (Linnaeus, 1758). Colors correspond to agricultural systems: Conservation Agriculture (CA, red), Conventional Agriculture (CVA, blue), Organic Agriculture (OA, green), and Organic Conservation Agriculture (OCA, yellow). Axes represent the constrained canonical axes, with values indicating the proportion of total variance explained by each axis. Asterisks denote statistically significant: * for *p-value* < 0.05 and ** for *p-value* < 0.01. Black arrows represent quantitative predictor variables of agricultural practices, with arrow length and direction indicating the strength and sign of their influence on community composition.
